# *APOE4* genotype and old age interact to impact cerebrovascular function, brain volume, and neuroinflammation in mice

**DOI:** 10.64898/2026.07.07.736860

**Authors:** Mackenzie N Kehmeier, Young D Choi, Abigail E Cullen, Benjamin Zimmerman, Thomas Leonhardt, Maddie Snyder, Tucker Cleveland, Naly Setthavongsack, Randall L. Woltjer, Martin M. Pike, Nabil J Alkayed, Ashley E Walker

**Author notes:** **Contact information**, Ashley E. Walker, PhD, Department of Human Physiology, University of Oregon, 139 Straub Hall, 1240 University of Oregon Eugene, Oregon, 97403, USA, https://vascularlab.uoregon.edu/. Authors contributed equally (Mackenzie N Kehmeier, Young D Choi).

## Abstract

Old age and the apolipoprotein E ε4 (*APOE4)* genotype are two of the greatest risk factors for late-onset Alzheimer’s disease (LOAD). However, the interaction between these is poorly understood, as most preclinical studies use young mice. Therefore, we assessed the interaction between *APOE* genotype and age across a comprehensive set of cerebrovascular and related outcomes. We performed *in vivo* imaging, *ex vivo* cerebral artery studies, behavioral tests, and molecular analyses in male and female homozygous *APOE3* and *APOE4* mice at ∼6 months (young) and ∼24 months (old). *APOE4* interacted with old age to lead to deficits in brain volume and greater microglia content. Old *APOE4* mice also exhibited greater cerebral artery vasoconstriction to endothelin-1 (ET-1) than old *APOE3* mice, a response concomitant with age-and genotype-related differences in the expression of ET-1 receptors and endothelin-converting enzyme. While we found several interactions between age and *APOE* genotype, only age impacted cognitive function, cerebral artery endothelial function, and arterial stiffness. In summary, we found that brain volume, neuroinflammation, and ET-1-related outcomes were influenced by the interaction of *APOE* genotype and age, while other outcomes were affected only by age. As such, an altered ET-1 response and greater neuroinflammation may contribute to the increased risk for LOAD in *APOE4* carriers.

## BACKGROUND

Historically, Alzheimer’s disease (AD) research in rodents focused on the damage caused by amyloid-β (Aβ).^1^ These studies ignored the primary risk factors for late-onset AD (LOAD): *old age*.^2^ Furthermore, the apolipoprotein E ε*4* (*APOE4*) genotype is the strongest genetic risk factor for LOAD, increasing disease risk by 2-13-fold compared with the neutral *APOE* ε*3* (*APOE3*) genotype.^3^ While the effects of old age and *APOE4* have been studied independently, very few studies have empirically examined their interaction, such as by preclinical studies conducted in models beyond middle age (18+ months in mice).^4,5^ As such, it is unclear how the interaction of age and *APOE* genotype impacts neurodegeneration, neuroinflammation, cognitive function, and cerebrovascular outcomes. Understanding how these risk factors interact is crucial to a better understanding of LOAD.

Reductions in cerebral blood flow and neurovascular coupling are potential contributors to the increased risk of LOAD. Indeed, dysregulation of cerebral blood flow is among the earliest biomarkers to change in the disease process.^6^ Independently, old age and the *APOE4* genotype are associated with impaired cerebral blood flow, limiting the delivery of key nutrients and oxygen to the brain, potentially serving as antecedents of neuroinflammation and neurodegeneration.^7,8^ Contributing to the reduced blood flow are impaired cerebral endothelial cells, key regulators of vasodilatory signaling.^9^ While impaired cerebral artery endothelium-dependent vasodilation is well established in old wildtype mice,^10^ it is unknown if the *APOE4* genotype influences this age-related response. In addition to vasodilatory signaling, vasoconstrictors also play a role in LOAD.

The endothelin-1 (ET-1) pathway not only has vasoactive effects but is also implicated in AD pathology. ET-1 is a potent vasoconstrictor of cerebral arteries and arterioles,^6^ and it has been shown that Aβ-induced vasoconstriction of capillaries is mediated through ET-1.^11^ Indeed, ET-1 is elevated in the brain with advancing age and in patients with AD.^12^ ET-1 also contributes to the greater vascular tone observed in older adults,^13^ while blockade of ET-1 signaling with

ET-1 receptor antagonists preserves arterial vasodilatory function in an AD mouse model.^14^ ET-1 is produced from the cleavage of Big ET-1 by endothelin converting enzymes (ECEs). ECEs not only cleave Big ET-1, but also degrade Aβ, as demonstrated by greater Aβ accumulation in ECE knockout AD mice.^15^ Thus, ET-1 and ECEs impact AD through multiple mechanisms; however, the interactive effects of old age and *APOE4* on cerebral blood flow, vasodilator/vasoconstrictor signaling, and ET-1 are unknown.

Therefore, the purpose of this study was to investigate the effects of age and the *APOE* genotype on brain volume, neuroinflammation, cognitive function, cerebral blood flow, and responses to vasodilators and vasoconstrictors. We hypothesized that there would be an interaction of old age and the *APOE4* genotype on these outcomes, such that old *APOE4* mice would have the greatest deficits compared with young *APOE4* and old *APOE3* mice.

Additionally, we investigated the impact on large elastic artery and cerebral artery stiffness, outcomes that are associated with LOAD and cognitive decline.^16,17^ Lastly, we assessed indirect calorimetry given the known effects of age and *APOE* genotype on these metabolic processes and fuel use. To perform this study, we compared young (6-month) and old (24-month) mice that were homozygous for *APOE3* or *APOE4*. Given known sex differences in AD risk, we also examined the interaction with sex for all outcomes.

## METHODS

### Animals

We purchased *APOE3* (JAX #029018) and *APOE4* (JAX #027894) mice from The Jackson Laboratories to establish a breeding colony. We studied male and female homozygous *APOE3* and *APOE4* mice at ∼6 months (young) and ∼24 months (old). Mice were fed a standard chow diet (Lab Diet, PicoLab Rodent Diet 20, 5053). Food and water were provided ad libitum, and mice were housed on a 12/12-hour light-dark cycle at 24°C. Mice were euthanized by exsanguination under isoflurane immediately prior to vascular studies. For immunohistochemistry tissue collection, mice were euthanized by thoracotomy followed by perfusion with heparinized saline under isoflurane anesthesia, and subsequent perfusion with 4% paraformaldehyde. All animal procedures conform to the Guide for the Care and Use of Laboratory Animals (8th edition, revised 2011) and were approved by the Institutional Animal Care and Use Committee at the University of Oregon.

### Magnetic resonance imaging (MRI)

MRI was performed at the OHSU Advanced Imaging Research Center using a Bruker-Biospin 11.75 T small animal MR system with a ParaVision 6.0 software platform, 10 cm inner diameter gradient set with a 72 mm (ID) and 60 mm (length) RF resonator for transmitting and an actively decoupled mouse head surface coil for receiving.^18^ Mice were anesthetized with a ketamine/xylazine mixture (1.0 mg xylazine/7 mg ketamine/100 g) in combination with low isoflurane (0.75%) in 100% oxygen. The mice were positioned with heads immobilized on an animal cradle. Body temperature of the mice was monitored and maintained at 37°C, and respiration was monitored (SA Instruments, Stony Brook, NY, United States). For each mouse, a coronal 25-slice T2-weighted image was acquired (ParaVision spin echo RARE, 256 × 256 matrix, 125 µm in-plane resolution, 0.5 mm slice width, TR 4000 ms, TE effective 23.64 ms,

RARE factor 8, 2 averages). These T2-weighted anatomical scans were used for positioning the blood flow image slice at a consistent position approximately 1.75 mm anterior to the anterior commissure. Cerebral blood flow (ml/min/100 g) was measured using arterial spin labeling (ASL), employing the flow-sensitive alternating inversion recovery rapid acquisition with relaxation enhancement pulse sequence (ParaVision FAIR-RARE), with TE/TR = 45.2/10000 ms, slice thickness =1 mm, number of slices = 1, matrix = 128 × 128, 250 µm in-plane resolution RARE factor = 72, and 23 turbo inversion recovery values ranging from 40 to 4400 ms, and acquisition time of 15 min. This sequence labels the inflowing blood by global inversion of the equilibrium magnetization.^19^ Cerebral blood flow maps (ml/100 g-min) were generated using the Bruker Paravision ASL perfusion processing macro and exported into JIM 9 software (Xinapse Systems LTD, Northants, United Kingdom) for further processing. Outlier values in brain pixels (outside 2 SDs) representing large arteries with high, pulsatile flow were excluded, thereby arriving at flows that consistently represent tissue microvascular blood flow. The identical FOV geometry offsets of the T2 and ASL images enabled the ROIs drawn on the T2 image to be readily overlaid onto the corresponding blood flow map for quantification. Mean blood flow was quantified for the whole brain, cortex, hippocampus, and thalamus, each defined anatomically using the corresponding T2 images. ASL map voxels containing ventricular contributions were excluded. The Allen Brain Atlas was a guide to identify ventricles, hippocampus, cortex, and thalamus.

We implemented a custom preprocessing pipeline using FSL (v6.0) and ANTs (v2.4.4) to analyze mouse brain anatomy from high-resolution T2-weighted MRI data.^18^ T2-weighted volumes were first bias-field corrected using *N4BiasFieldCorrection* to mitigate intensity inhomogeneities. Brain extraction was performed using *bet4animal*. Each subject’s brain-extracted image was registered to a mouse anatomical reference atlas. Registration was carried out using *antsRegistration* (ANTs v2.4.4) with a multistage pipeline consisting of rigid, affine, and SyN nonlinear transformations. Each step used mutual information and cross-correlation metrics with multi-resolution smoothing and shrink schedules to optimize spatial alignment. The resulting transformations were applied to the Badhwar atlas label map using *antsApplyTransforms.* Quality control images and Jacobian determinant maps were generated for all registrations to verify transformation accuracy and assess local volumetric distortions. Regional masks were derived from the transformed atlas label map to quantify volumetric measures across key anatomical structures, including the hippocampus, cortex, thalamus, hypothalamus, amygdala, striatum, corpus callosum, and ventricular system. Bilateral regions were merged to form composite masks where appropriate. Ventricular masks (including lateral, third, and fourth ventricles) were combined and subtracted from the whole-brain mask to derive a parenchymal (non-ventricular) brain volume. Regional and whole-brain volumes were quantified using *fslstats*, with voxel counts multiplied by voxel size to yield volume in mm³. All extracted measures were summarized in CSV format for statistical analysis.

### Cognitive function, motor coordination, and frailty

Learning and spatial memory were assessed via the Morris Water Maze. Mice underwent a 3-day training period, during which they performed 3 trials per session, 2 sessions per day. On the fourth day, a probe trial was performed, during which the platform was removed, and the mice swam in the arena for 60 seconds. All sessions were analyzed and recorded on EthoVision XT12 software (Noldus, Wageningen, the Netherlands).^20^

Nest building is a test of instinctual behavior and was performed over a 12-hour period. Mice were singly housed overnight in a fresh cage containing a condensed cotton nestlet. The following morning, the researcher scored the nests. Nests were scored on a scale of 0-5.^21^

Motor coordination testing was performed using a rotarod (47650 Rota-Rod NG, Ugo Basile, Gemonio, Italy) over a two-day period. The first day was a training session in which mice were required to stay on the rod rotating at 4 rpm for 90 seconds. The second day consisted of three trials, with a 10-minute rest between trials, during which the rod accelerated from 4 to 40 rpm, with a cut-off time of 5 minutes. The time was recorded for when the mouse fell from the rod or rotated around the rod two consecutive times.^22^

Anxiety was measured via open field testing. Mice were placed in a 40cm^2^ square arena with a white matte floor and walls. Mice were in the arena for 10 minutes, and their exploration was recorded using Ethovision XT12 software. The center of the 20×20 cm square was marked in the software, and the exploration criterion was defined as the amount of time the center of the mouse’s body was within the square. Mice that spent more time exploring have lower levels of anxiety.^20^

Mouse frailty was assessed using a previously studied 31-item frailty index (FI) based on established clinical signs of deterioration in C57BL/6J mice.^23^ FI assessments included evaluation of the integumentary, musculoskeletal, vestibulocochlear/auditory, ocular, nasal, digestive, urogenital, and respiratory systems, as well as signs of discomfort, body weight, and body surface temperature. The severity of each deficit was assessed and assigned a score of 0, 0.5, or 1, with higher scores indicating more severe frailty. Each mouse was examined at approximately the same time in the day, following the same order of assessments.

### Indirect Calorimetry

Mice were singly housed in an environmental cabinet for 5 days. The first 24 hours were excluded from the data analysis to account for the acclimatization period. Using metabolic cages (Promethion, Sable Systems, Las Vegas NV), we measured food and water intake, body weight, oxygen consumption (VO_2_), carbon dioxide production (VCO_2_), and water vapor during the 12:12-h light-dark cycles. Metascreen software (Sable Systems Inc.) was used for data collection. Data were then processed with ExpeData and the Macro Interpreter (Sable Systems Inc.) and analyzed hourly. VO_2_, VCO_2_ production, and expenditure (kcals/hour) were normalized to body mass.

### Vascular Reactivity

Endothelium-dependent vasodilation was assessed in ex vivo isolated pressurized posterior cerebral arteries (PCAs), as described in detail previously.^24,25^ Arteries were excised and cannulated onto glass micropipettes in a myograph chamber (Danish MyoTechnology Inc.) filled with a physiological salt solution. All the arteries were pre-constricted with phenylephrine (20-60 μM) to obtain 20-40% pre-constriction of the starting luminal diameter. Increases in luminal diameter in response to increasing concentrations of endothelium-dependent dilators acetylcholine (ACh: 1×10^-9^ to 1×10^-4^ M) and insulin (1 to 10000 ng/mL) were determined. After ACh and insulin responses, arteries were incubated for 30 minutes with 0.1 mM of Nω-Nitro-L-arginine methyl ester hydrochloride (L-NAME). ACh and insulin responses were repeated in the presence of LNAME to determine the contribution of nitric oxide synthases to endothelium-dependent dilation. Endothelium-independent dilation was measured by the increase in luminal diameter to sodium nitroprusside (SNP, 1×10^-10^ to 1×10^-4^ M). PCA vasoconstriction was also measured in response to potassium chloride (KCl: 10 to 80 mM) and ET-1 (1×10^-11^ to 1×10^-7^ M).

### Arterial stiffness

Passive arterial stiffness was measured ex vivo in the PCAs by the changes to lumen diameter and medial wall thickness in response to increasing intraluminal pressure.^26^ Passive stiffness was measured following a 60-minute incubation in a calcium-free solution to eliminate myogenic tone. Measurements were recorded in 5 cmH_2_O increments from 5 to 100 cmH_2_O (3.7 to 73.5 mmHg). From these data, stress-strain curves were created and β stiffness was calculated as previously described.^27^

Aortic stiffness was measured in vivo by pulse wave velocity.^24^ Mice were anesthetized using isoflurane (2% isoflurane in 100% oxygen) and placed supine on a temperature-controlled heating pad (37°C). Electrodes were placed on the distal portions of the limbs to record the ECG, while two Doppler transducers were positioned on the aortic arch and the abdominal aorta. The distance between the transducers was recorded. Using the Doppler signal processing workstation program (DSPW; Indus Industries, Webster, TX, USA), the time for a pulse to travel from the arch to the abdominal aorta was analyzed, and the time was then divided by the distance between the transducers. Two researchers independently analyzed the files, and the average was calculated.^28^

### Immunohistochemistry

The right hemisphere was fixed in 4% paraformaldehyde in PBS for 48 hours. Samples were transferred to a 100% PBS solution and stored at 4°C until further use. Brains were embedded in paraffin before being cut at 5 um thickness. Immunohistochemical staining was conducted using primary antibodies glial fibrillary acidic protein (GFAP, 1:500, Proteintech, Cat: 16825-1-AP) and ionized calcium-binding adaptor molecule 1 (Iba1, 1:5000, Proteintech, Cat: 10904-1-AP). Results were visualized after application of appropriate secondary antibodies using diaminobenzidine (DAB). Regions of the entorhinal cortex, hippocampus, and thalamus from these sections were imaged at 20x using a Leica Microscope (Leica, Wetzlar, Germany) and Leica LAS software. The percent positive area was analyzed in FIJI (ImageJ).

### Protein expression

The hippocampus and aorta were excised and frozen in liquid nitrogen. Samples were sonicated in total protein extraction reagent buffer with protease inhibitor cocktail for 10 seconds, three times at 70% power. Lysate protein concentration was assessed using a commercially available kit (Pierce BCA assay kit; Thermo Fisher, Waltham, MA). 4% Bio-Rad stain-free TGX gels were used for protein separation. The stain was activated and transferred onto a polyvinylidene difluoride transfer membrane. Membranes were imaged after transfer to ensure total protein transfer. Following imaging, membranes were blocked with 1.5% milk and then incubated in primary antibodies overnight in PBS containing 3% bovine serum albumin (BSA) and 0.05% sodium azide. Primary antibody concentrations were, respectively: APOE, 1:1,000 (ab52607, Abcam); ET-1, 1:500 (ab52607; ThermoFisher); ET_A_ receptor (ET_A_R), 1:500 (ab52607; ThermoFisher); ET_B_ receptor (ET_B_R), 1:1,000 (20964-1-AP; Proteintech). Following three washes for 10 minutes each with TBS-Tween (0.9% NaCl, 10 mM Tris, 0.1% Tween-20, pH 7.4), the membranes were incubated with a species-specific secondary antibody (Mouse: 7076, Cell Signaling; Rabbit: 7074, Cell Signaling) for 45 minutes. After secondary incubation, membranes were washed three times for 10 minutes each. Following washes, membranes were incubated with enhanced chemiluminescence (ECL) and imaged with a ChemiDoc Imaging System (Bio-Rad, Hercules, CA). Bio-Rad Image Lab was used to determine protein band density, and relative protein expression was calculated by normalizing protein density to stain-free blots. Values were normalized to the young *APOE3* male group.

### Gene expression

In samples of the cerebral cortex, mRNA gene expression was quantified for interleukin 1β (*Il1b*), NADPH oxidase 2 (*Nox2*), superoxide dismutase (*Sod1*)*, Sod2,* and *Sod3*. In samples of cerebral arteries (combined middle cerebral artery, posterior cerebral artery, and basilar artery), mRNA gene expression was quantified for ET-1 (*Edn1*)*, Ece1, Ece2,* and BK_ca_ channel (*Kcnma1*). RNA from the cortex and cerebral arteries was isolated by a standard protocol using Qiazol and RNAeasy Mini and Micro Kits (Qiagen, Hilden, Germany). Isolated RNA was quantified using a NanoDrop 2000 (Thermo Scientific, Waltham, MA), with further reverse transcription performed to produce cDNA with a Qiagen QuantiTect Reverse Transcription Kit. The cDNA samples underwent real-time qPCR with ThermoFisher PowerUp Sybr Green using a QuantStudio 5 real-time PCR system (ThermoFisher Scientific, Waltham, MA). mRNA expression was calculated using the 2^-ΔΔCT^ method, and 18S mRNA was used as a housekeeping control, with values normalized to the young *APOE3* male group.LJPrimers sequences are provided in **Supplemental Table 1.**

### Statistical analysis

Statistical analyses were performed with IBM SPSS (Version 26, Armonk, NY) and GraphPad Prism 10. A three-way analysis of variance (ANOVA) was used to determine the interaction effect and main effects of genotype, age, and sex. In the case of a significant F-value, post-hoc analyses were performed using a Tukey correction for preplanned comparisons. A repeated-measures ANOVA was used to assess group differences in dose responses and Morris Water Maze training trials. All data were tested for normality using a Shapiro-Wilk test. When data were not normally distributed, transformation was performed as noted in the figure legends. Significance was set at p<0.05, and values are represented as mean ± SE. Outliers were identified as those with z-scores>2 and removed from the dataset.

## RESULTS

### Animal Characteristics

Body mass was significantly greater in the old mice than in the young mice (young *APOE3*: 28.1±5.6 g, young *APOE4*: 28.3±4.7 g, old *APOE3*: 34.0±4.4 g, old *APOE4*: 35.8±9.3 g; main effect of age: p<0.0001). Old mice also had significantly greater heart mass, spleen mass, and white adipose tissue mass compared with the young mice, independent of *APOE* genotype (**Supplemental Table 2**). Frailty index was only assessed in old mice and did not differ between *APOE3* and *APOE4* mice (**Supplemental Table 2**).

### APOE4 is associated with lower brain volumes

Brain volumes were assessed from high-resolution T2-weighted MRI performed in live mice. A representative diagram of the registered brain regions is shown in **Fig 1A**. We found that *APOE4* mice had lower volumes than *APOE3* mice across the whole brain parenchyma, cerebral cortex, hippocampus, corpus callosum, hypothalamus, and striatum (**Fig 1B-D**, **Supplemental Fig 1**), while there was no effect of genotype in the amygdala and thalamus (**Supplemental Fig 1**). Furthermore, there was an interaction between genotype and age in the whole-brain parenchyma, such that old *APOE4* mice had the lowest volume (**Fig 1B**).

**Figure 1.**
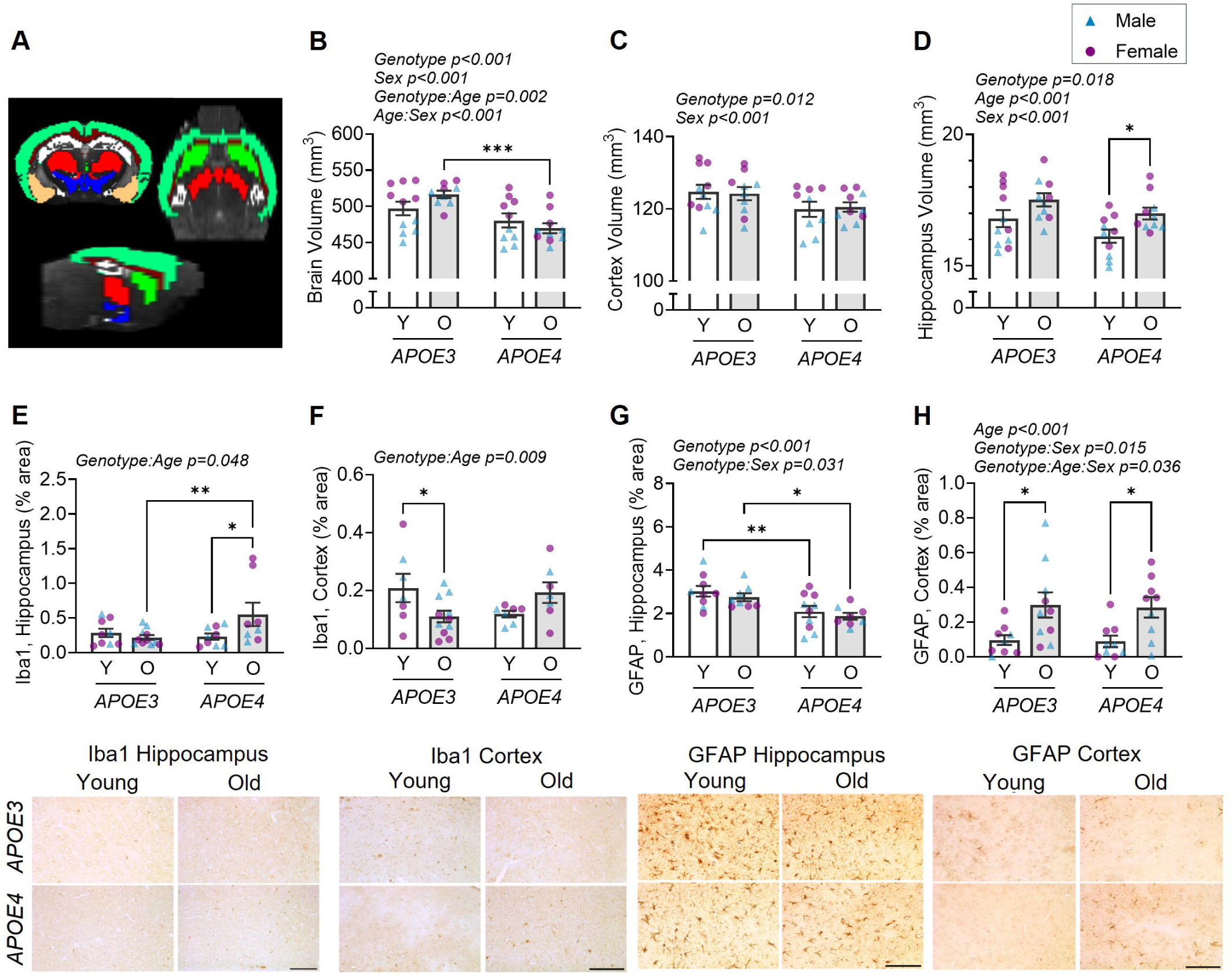
Brain volume is lower in *APOE4* mice, and *APOE4* interacts with old age to lead to greater microglia content, while the effects on astrocyte content are region dependent. From *in vivo* T2-weighted MRIs, (**A**) Representative image of brain volume analysis. Regions of interest: light green, cortex; white, hippocampus; beige, amygdala; brown, corpus callosum; red, thalamus; blue, hypothalamus; bright green, striatum. (**B**) Whole brain parenchyma volume, (**C**) cerebral cortex volume, and (**D**) hippocampus volume. For all regions, volumes were lower in *APOE4* mice compared with *APOE3* mice, but there was a variable impact of age and sex. Assessed by immunohistochemistry, there was an interaction of genotype and age for the content of Iba1 (microglia marker) in the (**E**) hippocampus and (**F**) cerebral cortex. For the content of GFAP (astrocyte marker), there was less in *APOE4* mice in the (**G**) hippocampus, while there was a greater content in older mice in the (**H**) cerebral cortex. Representative images are shown below (scale bar = 100 μm). Data analyzed by a three-way ANOVA with Tukey’s multiple comparisons. Y, young (∼6 months), O, old (∼24 months). *p<0.05, **p<0.01, ***p<0.001. n=3-6/group (genotype x age x sex). Data are mean ± SEM.

Counterintuitively, the hippocampus, thalamus, hypothalamus, and striatum were larger in old mice compared with young mice (**Fig 1D**, **Supplemental Fig 1**). Larger brain volumes in 6-month vs. 24-month mice align with the continued brain development in mice well beyond what is considered a young adult.^29^ As expected, the ventricles were larger in the old mice (**Supplemental Fig 1).** There was also a main effect of sex on the volumes of many brain regions, with females having larger volumes than males, but there were no interactions between sex and genotype. In summary, these data suggest that *APOE4* is associated with lower brain volumes, which are exacerbated by old age.

### APOE4 exacerbates age-related increases in neuroinflammatory markers

When assessing activated microglia content using Iba1, we found an interaction between age and genotype for Iba1-positive cells in the cortex and hippocampus. Specifically, while Iba1 was lower in *APOE3* with old age, it was higher in *APOE4* with old age (**Fig 1E-F**). Astrocyte content, assessed by GFAP, was lower in the cortex of *APOE4* mice, regardless of age (**Fig 1G**). In contrast, hippocampal GFAP-positive staining was higher in old mice compared with young mice, regardless of genotype (**Fig 1H**). As such, *APOE4* appears to promote an age-related accumulation of microglia, but not astrocytes.

We also assessed gene expression of inflammatory and pro- and anti-oxidant factors in the cerebral cortex. Old mice had a greater gene expression of NOX2 and IL-1β, independent of genotype (**Fig 2A,B**). There was an effect of sex (higher in males) in SOD1 and SOD3 expression, but not effects of age or genotype (**Fig 2C,E**). In contrast, *APOE4* mice had lower SOD2 expression compared with *APOE3* mice, regardless of age (**Fig 2D**). These results suggest that *APOE4* mice may have insufficient antioxidants to mitigate the age-related increases in pro-oxidant and pro-inflammatory factors.

**Figure 2.**
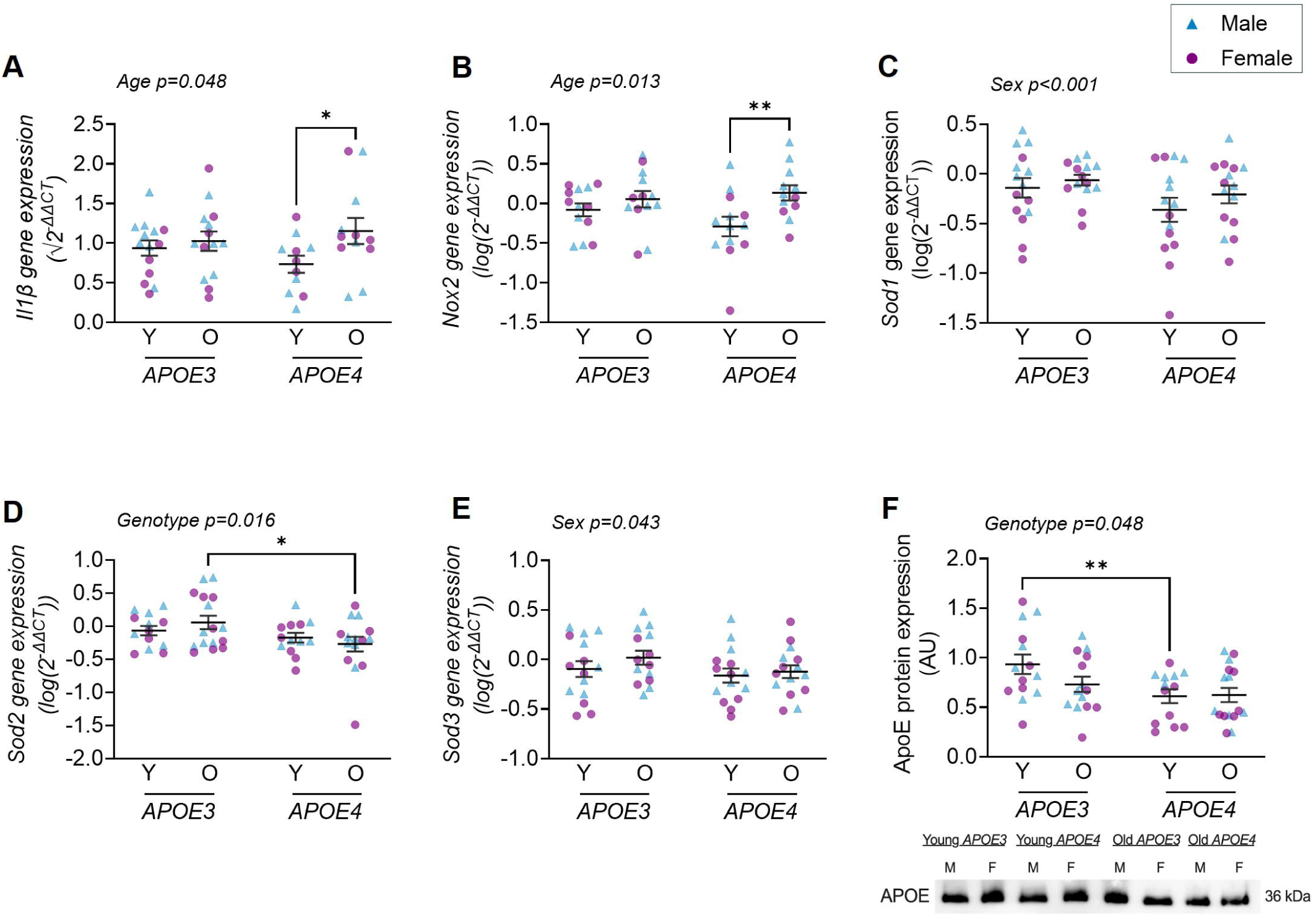
Pro-oxidant and pro-inflammatory gene expression is greater with old age, while antioxidant *Sod2* gene expression is lower in *APOE4* mice. Cerebral cortex gene expression of (**A**) interleukin 1β (*Il1b*) and (B) NADPH oxidase 2 (*Nox2*) were greater with age. (**C,E**) Superoxide dismutase 1 (*Sod1*) and 3 (*Sod3*) were lower in female mice, while (**D**) *Sod2* was lower in APOE4 mice. (**F**) Protein expression of ApoE in the hippocampus was lower in *APOE4* mice (data are normalized to a stain-free blot, **Supplemental** Figure 2). Data analyzed by a three-way ANOVA with Tukey’s multiple comparisons. Data for (A,F) were transformed by the square root, and data for (B,C,D,E) were transformed by log_10_. Y, young (∼6 months), O, old (∼24 months). *p<0.05, **p<0.01. n=4-8/group (genotype x age x sex). Data are mean ± SEM.

### APOE4 mice have lower ApoE protein expression

As in previous studies,^30^ we found that ApoE protein expression was lower in the hippocampus of young *APOE4* mice compared with young *APOE3* mice. There is a trend toward lower ApoE expression in old *APOE3* mice compared with young *APOE3* mice, and thus, we observed no differences in ApoE expression between the old mouse groups (**Fig 2F**).

### Cognitive function is impaired with old age, independent of APOE genotype

The Morris water maze was used to assess learning and spatial memory. There was a main effect of age on the training trials, with old mice performing worse than young mice regardless of genotype (**Fig 3A**). Old mice performed poorly on the probe trial compared with young mice, as shown by the young mice spending more time in the target vs. other quadrants, and this was independent of genotype (**Fig 3B**). Old mice had a slower swim velocity during the probe trial (**Fig 3C**). Similarly, old mice performed worse on the nest-building test and the accelerating rotarod, independent of *APOE* genotype (**Fig 3D,E**). During the open field test, old mice spent less time in the center of the arena, indicating greater anxiety, and moved a shorter distance during the test period, both of which were independent of *APOE* genotype (**Fig 3F,G**). As the old mice had impaired learning during the water maze training trials, motor impairments, and anxiety, the age-related decline in performance during the probe trial may not reflect memory impairment. Collectively, these results indicate that age has a strong impact on cognitive-related outcomes, including learning, instinctive behavior, motor control, and anxiety, and that the effects of age are much greater than those of *APOE* genotype in this context.

**Figure 3.**
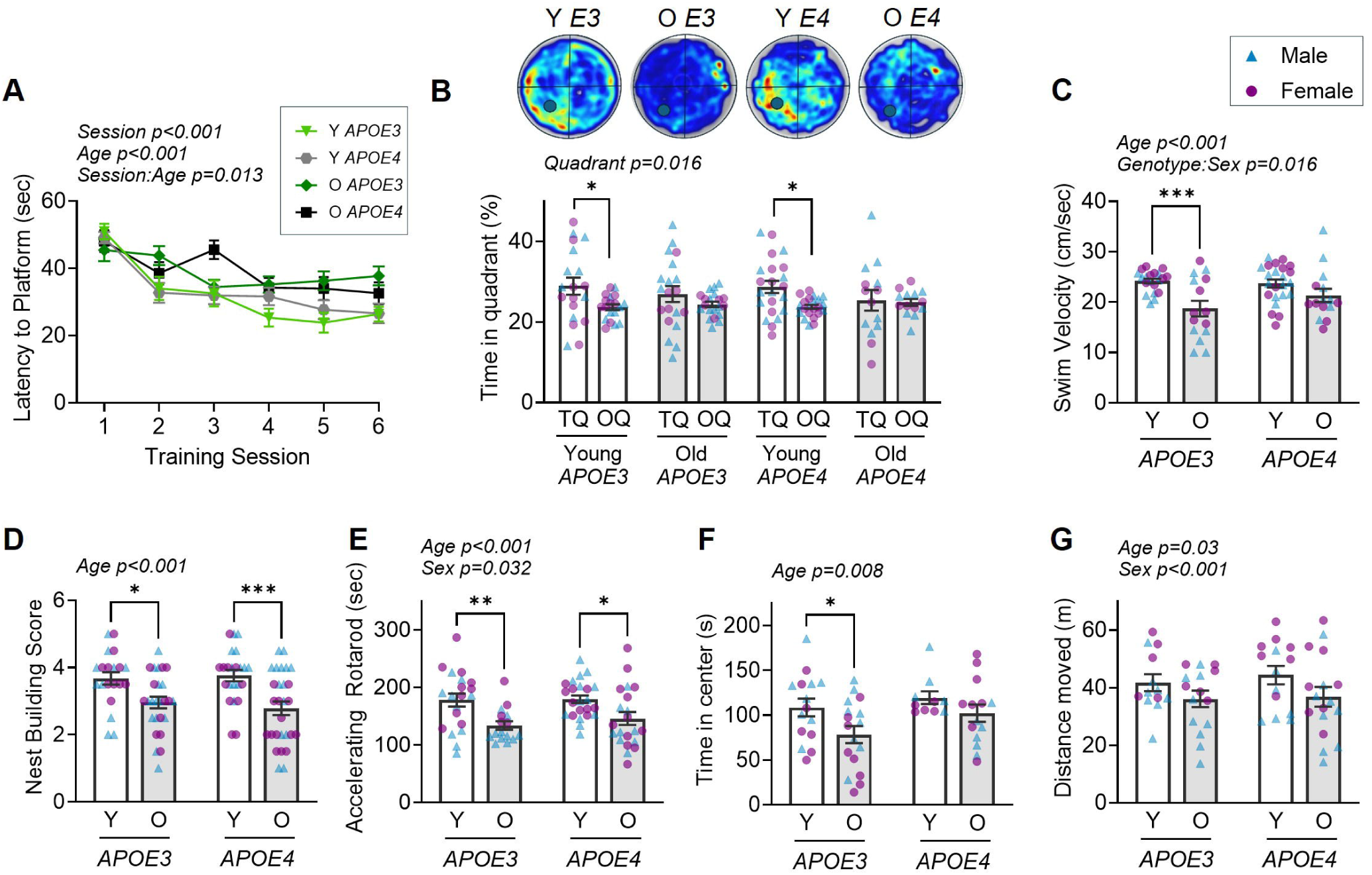
Cognitive function, motor coordination, and anxiety worsen with age but are not affected by *APOE* genotype. For the Morris Water Maze, (**A**) results from training trials were not different between groups, (**B**) time in target quadrant (TQ) vs. average of other quadrants (OQ) during probe trial was greater for the young groups, but not old groups, representative heat maps displayed above, and (**C**) swim velocity during the probe trial was lower in old groups. (**D**) Nest-building test and (**E**) accelerating rotarod results were impaired in the old groups but not affected by genotype. From the open field test, (**F**) time spent in the center of the arena and (**G**) total distance moved were lower in old groups but not affected by genotype. A three-way ANOVA with Tukey’s multiple comparisons was used for analysis, except for training trials, which were analyzed by a repeated measures ANOVA. Y, young (∼6 months), O, old (∼24 months). *p<0.05, **p<0.01, ***p<0.001. n=5-14/group (genotype x age x sex). Data are mean ± SEM.

### APOE4 impacts the age-related changes in vasoconstrictor responses and the ET-1 pathway

Previous studies have shown that ET-1 is greater in the brains of AD patients,^12,31^ and vasoconstriction to Aβ is mediated by the effects of ET-1.^32^ We found an interaction of age and *APOE* genotype on ET-1-induced vasoconstriction of the PCA. Specifically, the maximal ET-1-induced vasoconstriction was 18% lower in old *APOE3* mice compared with young *APOE3* mice (**Fig 4A**), and was similarly different for the dose-response (**Fig 4B**). These results are consistent with studies in wild-type C57BL/6 mice, which show that the response to ET-1 diminishes with age.^33^ In contrast, the PCAs from old *APOE4* mice did not differ in ET-1-induced vasoconstriction when compared with young *APOE4* mice (for maximal or dose responses, **Fig 4A,B**). As a result, there was greater ET-1-mediated PCA vasoconstriction in old *APOE4* mice compared with old *APOE3* mice. These results suggest that *APOE4* abrogates the age-related loss of ET-1 responsiveness observed in *APOE3* and wild-type mice.

**Figure 4.**
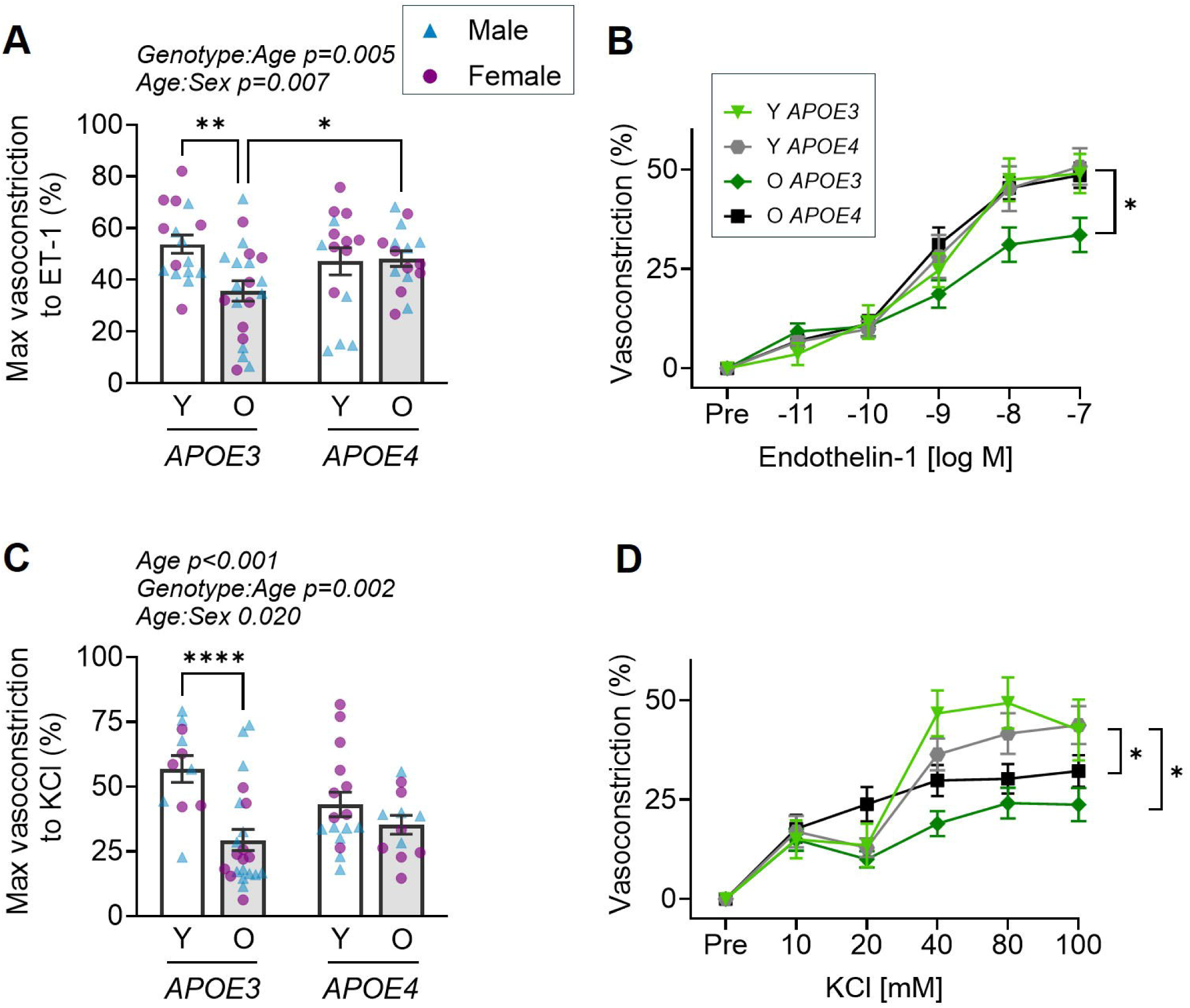
Cerebral arteries from old *APOE4* mice have a hyperactive vasoconstrictor response. In isolated posterior cerebral arteries, old *APOE4* had a greater (**A**) maximal or (**B**) dose-response vasoconstriction to endothelin-1 (ET-1) compared with old *APOE3* mice. The age-related decline in (**C**) maximal and (**D**) dose-response vasoconstriction to potassium chloride (KCl) was also absent in *APOE4* mice. A three-way ANOVA with Tukey’s multiple comparisons was used to analyze the maximal responses, and a repeated measures ANOVA was used to analyze the dose responses. Y, young (∼6 months), O, old (∼24 months). *p<0.05, **p<0.01, ****p<0.0001. n= 5-12/group (genotype x age x sex), data are mean ± SEM

To determine if the *APOE* effects on ET-1 vasoconstriction were a failure of vasoconstrictor signaling, we also tested cerebral artery vasoconstriction in response to KCl. There was an interaction between age and genotype in the vasoconstriction to KCL, such that there were age-related reductions in the response in *APOE3* mice but not in *APOE4* mice (**Fig 4C,D**). However, unlike ET-1, there was no difference in KCl-induced vasoconstriction between old *APOE3* and *APOE4* mice. Thus, while an interaction between *APOE* genotype and age was found for both ET-1 and KCl vasoconstriction, among old mice there was only an effect of genotype for ET-1.

To better understand the underlying cause of the lessened ET-1 responsiveness in old *APOE4*, we assessed the expression of ET-1-related genes and proteins. We found that aortic ET-1 protein expression was higher in *APOE4* mice than in *APOE3* mice and lower in old mice than in young mice (**Fig 5A**). ET_B_ receptors (ET_B_R) are primarily responsible for vasodilation via endothelial cells, an action that opposes ET_A_ receptor (ET_A_R)-mediated vasoconstriction in smooth muscle.^34^ We found an interaction of age and genotype for hippocampal ET_A_, such that ET_A_ protein expression was greater in young *APOE3* mice than young *APOE4* and old *APOE3* mice (**Fig 5B**). In contrast, ET_B_R expression was greater with age in the hippocampus for both genotypes (**Fig 5C**). These results provide initial insight into ET-1 signaling in the brain but are limited because they were not performed in vascular tissue, as such we also measured gene expression for ET-1 pathway components in cerebral arteries. There was an interaction between age and genotype in cerebral artery expression of pre-ET-1 (*Edn1*), with old *APOE4* mice exhibiting the highest expression (**Fig 5E**). ECEs catalyze the cleavage of Big ET-1 to active ET-1, but they are also known to cleave Aβ precursor protein (AβPP). *APOE4* mice had lower cerebral artery ECE1 expression, regardless of age (**Fig 5F**). For ECE2, there was an interaction between age and genotype, with lower expression in old vs. young *APOE3* mice, but no age-related differences in *APOE4* mice (**Fig 5G**). Lastly, we measured expression of the large-conductance calcium-activated K+ channel subunit α (*Kcnma1*), which was higher in the older groups, regardless of genotype (**Fig 5H**). In summary, in *APOE3* mice, ET-1-induced vasoconstriction declines with age, coinciding with lower ET_A_ expression and higher ET_B_R expression. However, the potential mechanisms underlying the sustained ET-induced vasoconstriction in old *APOE4* mice are less clear but may be mediated by preserved ET_A_ expression. There are also opposing results between ET-1 hippocampal protein expression and cerebral artery gene expression, with hippocampal expression higher in young *APOE4* and cerebral artery expression higher in old *APOE4*. Furthermore, the lower ECE1 expression in *APOE4* mice has implications for Aβ degradation.

**Figure 5.**
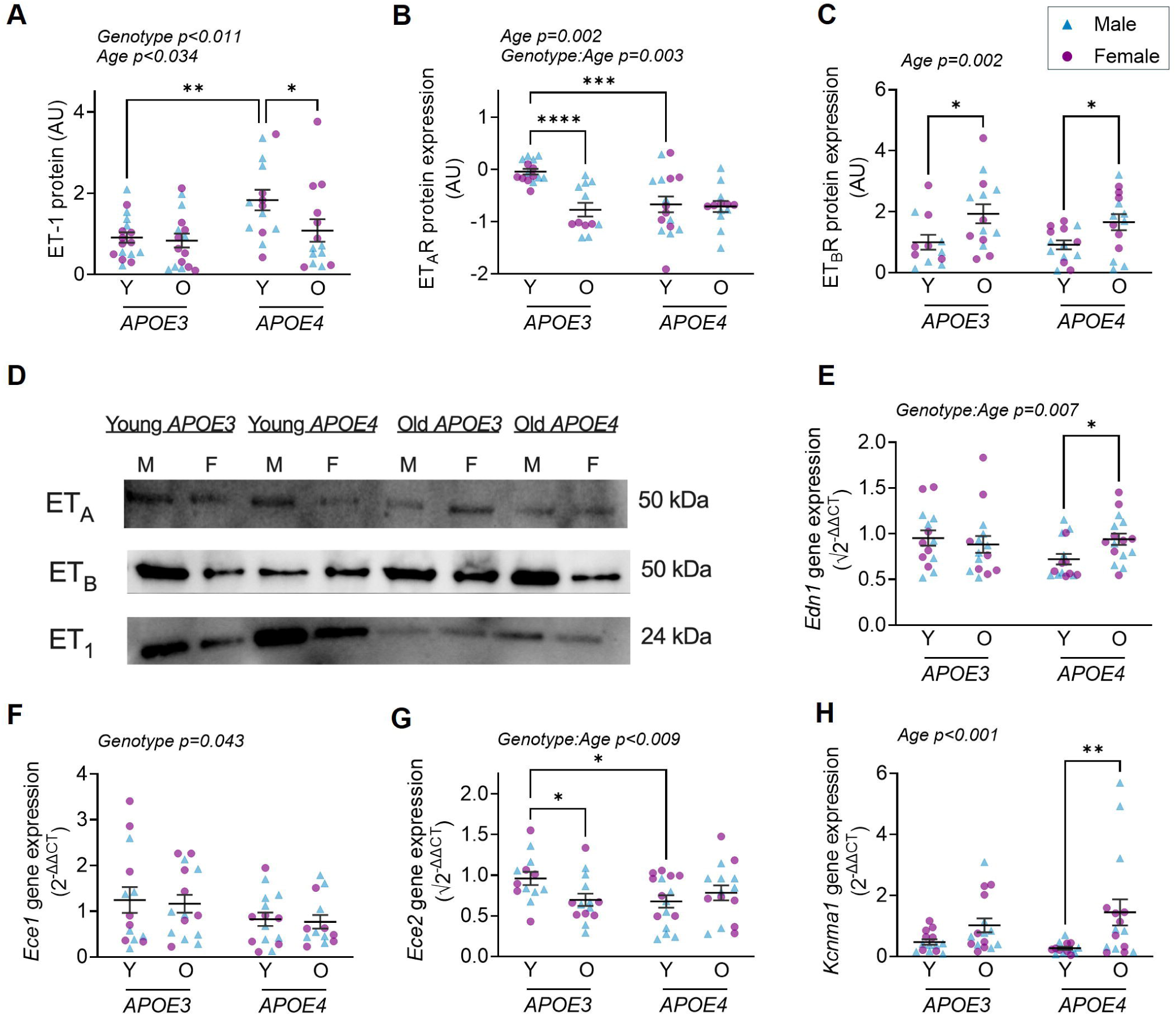
*APOE* genotype and age differentially affect expression of endothelin-1 and related pathway components. (**A**) Protein expression of endothelin-1 (ET-1) in the aorta was highest in the young *APOE4* mice, while expression of ET-1 receptors in the hippocampus (**B**) was highest in the young *APOE3* mice for ET_A_R, and (**C**) was greater with old age for ET_B_R. (**D**) Western blot representative images (data are normalized to stain-free blots, **Supplemental** Figure 2). Cerebral artery gene expression of (**E**) ET-1 (*Edn1*) was greater with aging in *APOE4* mice, while expression of (**F**) ET-1 converting enzyme 1 (*Ece1*) was lower in *APOE4* and (**G**) *Ece2* was lower with age in *APOE3* but not *APOE4* mice. (**H**) Cerebral artery gene expression of BK_ca_ channel (*Kcnma1*) was greater with age. Data analyzed by a three-way ANOVA with Tukey’s multiple comparisons. Data for (B) were transformed by log_10_ and data for (E,G) were transformed by square root. Y, young (∼6 months), O, old (∼24 months). *p<0.05, **p<0.01. n=5-9/group (genotype x age x sex). Data are mean ± SEM.

### Cerebral artery endothelium-dependent vasodilation is dependent on age but not APOE genotype

Given the effects of genotype and age on vasoconstrictor responses, we also sought to understand their influence on vasodilation. The PCAs from old mice had impaired endothelium-dependent vasodilation to ACh and insulin compared to the young groups, but there was no main effect of *APOE* genotype (**Fig 6A-D**). The vasodilation to ACh and insulin was entirely mediated by nitric oxide, as indicated by little to no vasodilation in the presence of the nitric oxide synthase inhibitor, LNAME (**Fig 6B,D**). Endothelium-independent vasodilation did not differ between groups, as assessed by the response to sodium nitroprusside, indicating that the differences in vasodilation were due to age-related impairment of endothelial cells (**Fig 6H**). Sensitivity (EC_50_) to vasodilators and the amount of preconstriction did not differ between groups (**Supplemental Table 3**). Overall, these studies indicated that age has a strong effect on endothelial function, while *APOE* genotype had little effect.

**Figure 6.**
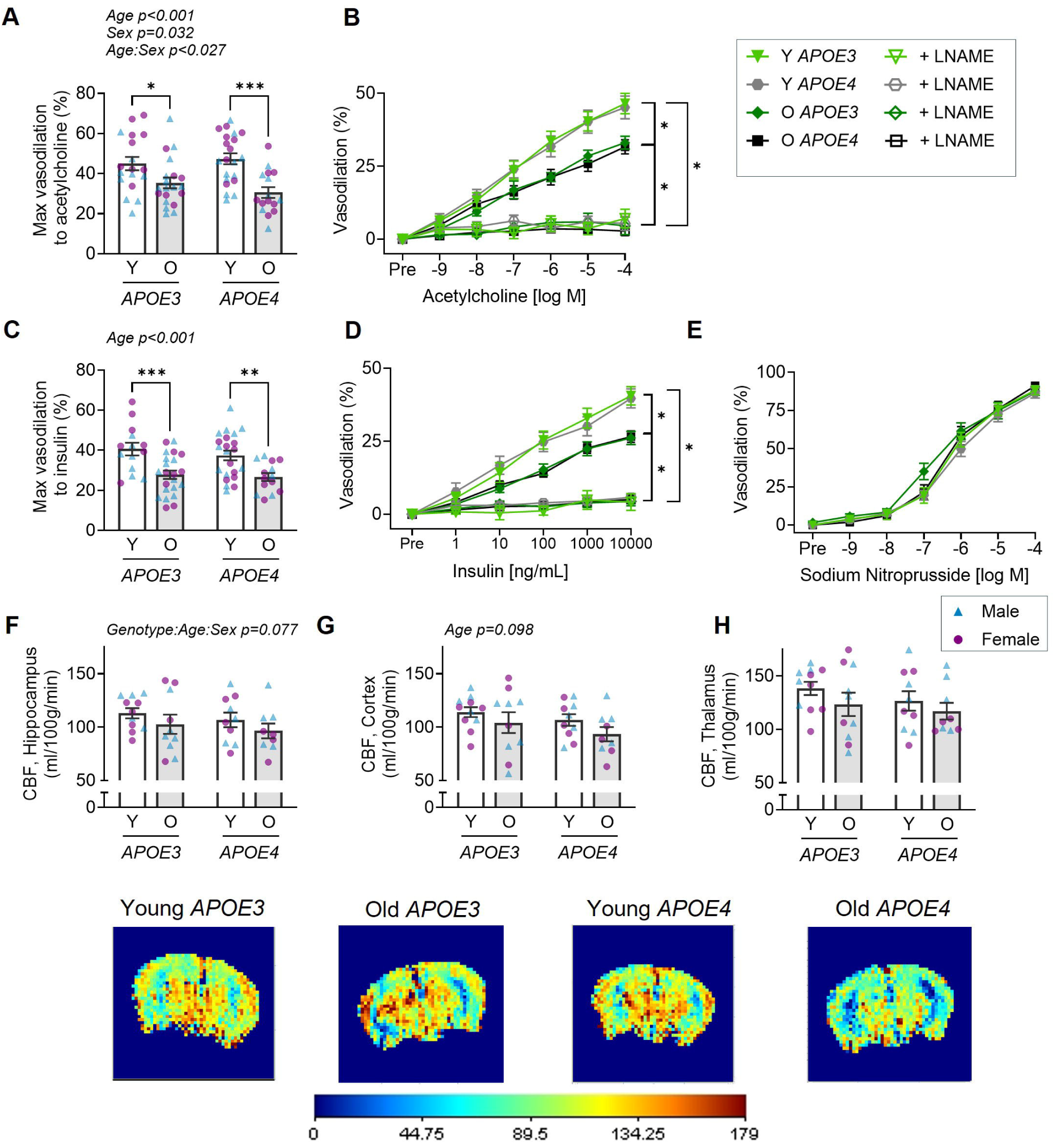
Endothelium-dependent vasodilation and cerebral blood flow were not affected by *APOE* genotype. In isolated posterior cerebral arteries, endothelium-dependent vasodilation to acetylcholine was measured, presented as (**A**) the maximal response and (**B**) the dose-response in the presence and absence of nitric oxide inhibitor, LNAME. Endothelium-dependent vasodilation was also measured in response to insulin, presented as (**C**) the maximal response and (**D**) the dose-response in the presence and absence of LNAME. The vasodilatory response to acetylcholine and insulin was impaired with age, but not affected by APOE genotype. (**E**) Endothelium-independent vasodilation in response to sodium nitroprusside was not different between groups. Using arterial spin labeling MRI, cerebral blood flow (CBF) was measured in the (**F**) hippocampus, (**G**) cortex, and (**H**) thalamus, and only non-statistically significant trends were observed. Representative images are shown below with a scale bar in ml/100g/min. A three-way ANOVA with Tukey’s multiple comparisons was used for analysis, except for dose responses, which were analyzed by repeated measures ANOVA. Y, young (∼6 months), O, old (∼24 months). *p<0.05, **p<0.01, ***p<0.001. n=4-13/group (genotype x age x sex). Data are mean ± SEM

### Effects of genotype, age, and sex on cerebral blood flow

Cerebral blood flow was measured by ASL MRI in the hippocampus, cortex, and thalamus. There was substantial variability in these outcomes, which limited our statistical power to detect differences. In the hippocampus, there was a trend towards an interaction of genotype, age, and sex on blood flow. Specifically, there was a trend toward lower hippocampal blood flow with age in *APOE3* males, but not in *APOE4* mice or female mice (**Fig 6F**). In the cortex, there was a trend for lower blood flow in old mice, regardless of genotype (**Fig 6G**). There was no influence of genotype, age, or sex on blood flow in the thalamus (**Fig 6H**).

### Cerebral artery and large elastic artery stiffness are greater with age, but are not influenced by APOE genotype

Greater arterial stiffness is highly correlated with impaired cerebral blood flow and cognitive decline.^35^ Thus, we assessed the stiffness of both cerebral and large elastic arteries. Passive stiffness was assessed in PCAs incubated in a solution free from Ca^2+^ to eliminate smooth muscle tone. Stress-strain curves were generated and used to calculate β stiffness (**Fig 7A**). Cerebral artery β stiffness was greater with age, independent of *APOE* genotype (**Fig 7A,B**). Aortic stiffening was measured in vivo by pulse wave velocity. As with the cerebral arteries, aortic stiffness was greater with age, regardless of *APOE* genotype (**Fig 7C**). These findings indicated that neither large elastic artery stiffness nor small cerebral artery stiffness is affected by *APOE* genotype.

**Figure 7.**
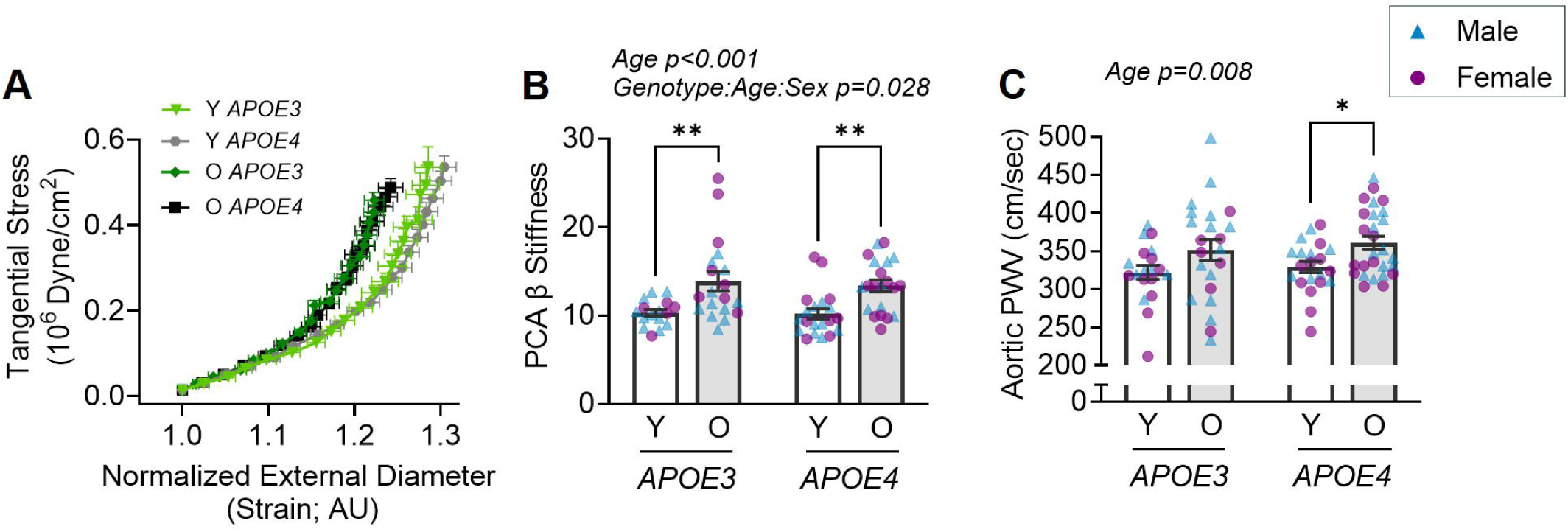
Arterial stiffness is greater in old age but is not affected by *APOE* genotype. In posterior cerebral arteries incubated in a calcium-free solution, the passive response to increasing pressure was used to generate (**A**) stress-strain curves and calculate (**B**) β-stiffness. (**C**) Aortic stiffness was measured by pulse wave velocity (PWV). Aortic and cerebral arterial stiffness were greater in old age but were not affected by *APOE* genotype. A three-way ANOVA with Tukey’s multiple comparisons test was used for the analysis. Y, young (∼6 months), O, old (∼24 months). *p<0.05, **p<0.01. n=5-14/group (genotype x age x sex). Data are mean ± SEM.

### Metabolism is influenced by age and APOE

In vivo whole-body metabolism was measured in a subset of mice via indirect calorimetry for four days. There was no statistically significant difference for light-cycle VO_2_ (**Fig 8A**). The old mice had a lower dark-cycle VO_2_ and light- and dark-cycle VCO_2_, regardless of genotype, compared with the young mice (**Fig 8B,D,E**). For light- and dark-cycle respiratory exchange ratio (RER), the old *APOE3* mice were lower than both the young *APOE3* mice and the old *APOE4* mice (**Fig 8C,F**). These results suggest that *APOE4* prevents the age-related declines in RER observed in *APOE3* mice.

**Figure 8.**
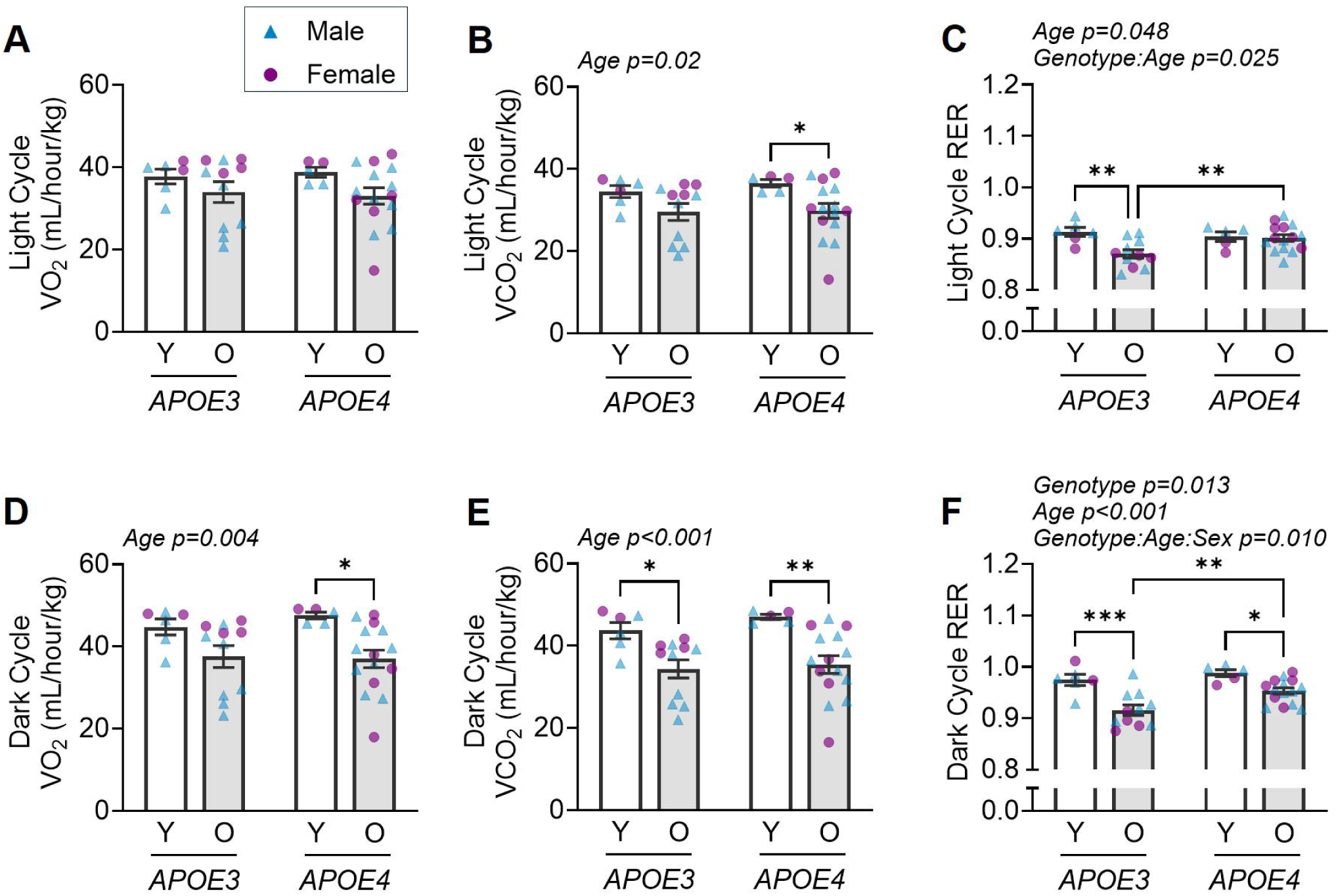
Age and *APOE* genotype impact whole-body metabolism. Using metabolic cages, metabolic markers were measured during the (**A-C**) light cycle and the (**D-F**) dark cycle, including (**A,D**) VO_2_, (**B,E**) VCO_2_, and (**C,F**) respiratory exchange ratio (RER). Old mice had lower light-cycle VCO_2_ and dark-cycle VO_2_ and VCO_2_, but there was no effect of genotype. Light-cycle RER was lowest in the old *APOE3* mice, while dark-cycle RER was affected by age, genotype, and the interaction of age, genotype, and sex. A three-way ANOVA with Tukey’s multiple comparisons was used for analysis. Y, young (∼6 months), O, old (∼24 months). *p<0.05, **p<0.01, ***p<0.001. n=2-9/group. Data are mean ± SEM.

## DISCUSSION

We assessed the interaction of age and *APOE* genotype on a comprehensive set of cerebrovascular and related outcomes. We found that *APOE4* mice exhibited more pronounced age-related deficits in brain volume, concomitant with greater microglia content. While age-related increases in pro-oxidant and pro-inflammatory gene expression occurred regardless of genotype, *APOE4* mice exhibited lower SOD2 antioxidant gene expression, suggesting a potential oxidative imbalance. We further found an age-related decline in ET-1-mediated cerebral artery vasoconstriction in *APOE3* mice but not in *APOE4* mice. This was concurrent with differences in ET-1-related protein and gene expression, which may suggest ET-1 dysregulation and could have implications for Aβ degradation in *APOE4* carriers. While we found several interactions between age and *APOE* genotype, numerous outcomes were influenced only by age. Old age led to impaired cognitive function, impaired endothelium-dependent vasodilation in cerebral arteries, and greater arterial stiffness, all of which were independent of *APOE* genotype. Thus, in this study, we found that brain volume, neuroinflammation, and ET-1-related outcomes were influenced by the interaction of *APOE* genotype and age, while other outcomes were affected only by age.

Studies in human subjects consistently find that *APOE4* carriers have lower brain volumes compared with non-carriers, even among cognitively healthy individuals.^36^ We found a similar result: *APOE4* mice had lower volumes in the cortex, hippocampus, corpus callosum, hypothalamus, and striatum compared with *APOE3* mice. However, we also unexpectedly found higher brain volumes in the old mice than in the young mice across many brain regions. This is consistent with a previous study suggesting that brain volumes continue to increase with age until ∼11 months in mice.^29^ Thus, despite our use of 6-month-old mice for our young group, it appears likely that brain development was still underway. A recent study has similarly found increases in brain volume with age but found no effect of *APOE* genotype in mice.^37^ However, the previous *APOE* study used ex vivo MRI, whereas our study was conducted in vivo, which may have allowed us to identify smaller genotypic differences. As such, *APOE4* mice appear to recapitulate the lower brain volumes observed in human *APOE4* carriers when measured by in vivo MRI.

Human *APOE4* carriers also have greater numbers of activated microglia^38^ and a heightened pro-inflammatory state.^39^ *APOE4* microglia also exhibit elevated secretion of pro-inflammatory cytokines.^40,41^ However, these activated *APOE4* microglia also have an impaired phagocytic function, leading to reduced Aβ clearance.^42^ In our study, we likewise found greater numbers of microglia in old *APOE4* mice, and, while not statistically significant, the age-related increases in pro-inflammatory gene expression trend toward greater in *APOE4* mice*. APOE4* is also associated with greater astrocyte activation, impaired astrocytic lipid transport, and reduced astrocytic end-feet contact with blood vessels, potentially leading to increased oxidative stress, impaired synaptic transmission, and increased blood-brain barrier permeability.^39,43,44^ Interestingly, we found that *APOE4*, regardless of age, was associated with fewer astrocytes in the hippocampus. Future studies of astrocyte function will be needed to determine whether this reduction in cell number has beneficial or detrimental effects.

Often coinciding with a pro-inflammatory environment is an increase in oxidative stress.^45^ In this study, we found that the expression of pro-oxidant NOX2 is greater with old age in cerebrovascular tissues, as previously shown.^27^ Interestingly, we found that antioxidant SOD2 expression was lower in cerebrovascular tissue from *APOE4* mice, consistent with our previous findings in young female *APOE4* mice.^46^ This is notable since SOD2 is specific to mitochondria, and *APOE4* is associated with mitochondrial impairment.^47,48^ As such, whereas age-related increases in pro-oxidants in *APOE3* mice may be counterbalanced by stronger antioxidant defenses, our findings suggest that *APOE4* shifts this balance toward heightened oxidative stress with advancing age.

Despite differences in brain volume and neuroinflammatory markers by genotype, we did not observe any cognitive impairment by genotype. This is inconsistent with known effects in humans and mice. A meta-analysis of cognitively healthy older adults indicates that *APOE4* carriers have worse cognitive performance than non-carriers, with larger differences emerging in old age.^49,50^ A more recent meta-analysis reports that *APOE4* mice have cognitive deficits irrespective of sex or age, depending on the experimental test condition.^49^ In this study, genotype-related cognitive impairments were likely overwhelmed by the effects of old age. Memory assessments in mice are confounded by differences in visual and motor function, two variables that worsen markedly with age. Thus, the discordance of our cognitive results with the brain volume and neuroinflammatory results is likely due to limitations of the behavioral tests used in the present study.

Age is generally associated with a loss of vasoconstrictor responsiveness. Older humans exhibit reduced ET-1-mediated vasoconstriction in the skeletal muscle circulation compared with young adults.^13^ We have similarly shown this in cerebral arteries of wildtype C57BL6 mice,^33^ and others have shown this in the coronary resistance arterioles of rats.^51^ In the present study, we found that old *APOE3* mice had the expected lower responsiveness to ET-1, which could be explained by the age-related decline in ET_A_R (primarily vasoconstrictor) and an elevation in ET_B_R (primarily vasodilator) expression. Compared with the old *APOE3* mice, the old *APOE4* mice were hypercontractile to ET-1. Interestingly, we recently found that WSB/EiJ mice, a strain more susceptible to cerebral amyloid angiopathy, also lack the age-related reductions in ET-1 responsiveness.^33^ The 5xFAD mouse also exhibits a hyperactive constriction phenotype,^52^ suggesting that this finding spans multiple AD-related models. In the 5xFAD mouse, this hypercontractile state was found to be mediated by impairment of BK_Ca_ channels.^52^ In our study, we find no differences in BK_Ca_ expression by *APOE* genotype, but it remains to be determined whether channel activity underlies the effects of *APOE4*. The lack of genotype differences in resting cerebral blood flow in our study could suggest that the response to vasoconstrictors does not affect perfusion. However, high variability in blood flow measures limited our ability to detect differences, and vasoconstrictor responsiveness may affect neurovascular coupling rather than global blood flow.

As important as the response to ET-1 is the amount present in the brain. ET-1 expression is elevated with advancing age and AD in the human brain.^12,53^ We found that ET-1 expression was not quantifiable in the hippocampus; however, in the aorta, ET-1 expression was higher in *APOE4* mice, at least at young ages. Furthermore, the pathway for producing ET-1 also has implications for AD pathology. Higher ECE expression may be detrimental, as it increases ET-1 production, but it may also be beneficial, as it increases Aβ breakdown.^54^ ECE1 gene expression is lower in cerebrovascular tissues from AD patients,^32^ and we similarly found that ECE1 gene expression was lower in *APOE4* cerebral arteries. This lower cerebrovascular ECE1 could decrease Aβ degradation, particularly near the vasculature, in *APOE4* individuals.

We found that *APOE4* mice had a higher dark cycle RER than *APOE3* mice, consistent with a preference for carbohydrate metabolism. The differences seen in RER are due to a slight increase in VCO_2_ and a slight decrease in VO_2_. In a previous study in 18-month-old male mice, there was no difference in RER between *APOE3* and *APOE4*.^55^ Our divergent findings could be due to sex, as we observed an interaction among genotype, age, and sex. A limitation of the metabolic studies is that we did not measure activity, and the differences in RER may reflect higher activity in the *APOE4* mice. Thus, our results suggest lower lipid utilization in *APOE4* mice, but the role of activity in these findings needs further investigation.

The results of this study are limited to the context in which these experiments were performed. Notably, the mice were fed a normal chow diet, which is low in fat and cholesterol. As a primary phenotype of *APOE4* is altered lipid transport,^56^ we may have observed greater differences between *APOE3* and *APOE4* mice had we fed them a high-fat diet. Furthermore, these mice lacked humanized *A*β*PP*. As *APOE4* plays a role in Aβ trafficking, higher Aβ levels could alter *APOE4* interactions with aging. One limitation in our study is that the normal chow diet contained soybean meal. Our previous study suggests that soybean meal (a phytoestrogen) can improve cerebrovascular and cognitive function.^24^ The effects of estrogen, and likely phytoestrogens, in the context of *APOE4* are complex, as we recently demonstrated that *APOE4* mice are resistant to the beneficial effects of estradiol on cerebral arteries.^46^ Thus, further studies are needed to understand the effects of diet and Aβ on the interaction between age and *APOE* genotype.

In conclusion, we found that the interaction of old age and the *APOE4* genotype leads to lower brain volumes, heightened neuroinflammation, and modulation of ET-1 signaling. However, there were several outcomes in which the effects of aging were independent of *APOE* genotype, including cerebral artery endothelial function, arterial stiffness, and cognitive function. These studies were performed in mice that did not express humanized AβPP and were fed a normal chow diet. As such, further studies are needed to understand how additional pathological or metabolic risk factors influence the interaction of *APOE4* and old age.

