## Supplemental tables and figures for "*APOE4* genotype and old age interact to impact cerebrovascular function, brain volume, and neuroinflammation in mice"

### DATA SUPPLEMENT

*Authors contributed equally

**Supplemental Table 1.** qPCR Primers

| **Primer** | **Forward** | **Reverse** |
|---|---|---|
| *Il1β* | GCAACTGTTCCTGAACTCAACT | ATCTTTTGGGGTCCGTCAACT |
| *Nox2* | TCCCAGAGAACACAGCATAAC | CTAGCCTGCTTATGGGATTCTT |
| *Sod1* | AACCAGTTGTGTTGTCAGGAC | CCACCATGTTTCTTAGAGTGAGG |
| *Sod2* | CAGACCTGCCTTACGACTATGG | CTCGGTGGCGTTGAGATTGTT |
| *Sod3* | CCTTCTTGTTCTACGGCTTGC | TCGCCTATCTTCTCAACCAGG |
| *Edn1* | TTTCCCGTGATCTTCTCTCTGC | CGCCTACCTGTTTCTGGAGC |
| *Ece1* | TCTCCGAGGGCGATGTGTA | CTTCTCCACCGAGGTCCGA |
| *Ece2* | GGGAAAACATCGCCGATAATGG | CGAAGTGCCGAAGGAAGTCTC |
| *Kcnma1* | TCACGGAACTCGCTAAGCC | AATGTGCGTCCCACTGTTTTT |

**Supplemental Table 2.** Animal Characteristics

| **Variable** | **Young *APOE3*** | **Young *APOE4*** | **Old *APOE*3** | **Old *APOE4*** |
|---|---|---|---|---|
| n | 20 | 24 | 26 | 22 |
| Age (months)[a] | 6.7 ± 0.2 | 6.5 ± 0.2 | 24 ± 0.4*† | 23 ± 0.9*† |
| Sex (%F) | 20 (45%) | 24 (46%) | 26 (50%) | 22 (50%) |
| Body mass (g)[a] | 28.1 ± 5.6 | 28.3 ± 4.7 | 34.0 ± 4.4*† | 35.8 ± 9.3*† |
| Heart mass (mg)[a] | 0.15 ± 0.03 | 0.15 ± 0.02 | 0.18 ± 0.03*† | 0.19 ± 0.03*† |
| Percent heart:body mass | 0.56 ± 0.1 | 0.52 ± 0.01 | 0.52 ± 0.1 | 0.55 ± 0.1 |
| Liver mass (mg)[a] | 1.53 ± 0.4 | 1.53 ± 0.4 | 1.82 ± 0.4 | 1.68 ± 0.7 |
| Percent liver:body mass | 5.39 ± 0.6 | 5.37 ± 0.7 | 5.39 ± 1.2 | 4.67 ± 1.4 |
| Spleen mass (mg)[a] | 0.1 ± 0.01 | 0.09 ± 0.01 | 0.16 ± 0.1† | 0.16 ± 0.1*† |
| Percent spleen:body mass | 0.38 ± 0.08 | 0.33 ± 0.07 | 0.48 ± 0.4 | 0.47 ± 0.3 |
| WAT mass (mg)[a] | 0.65 ± 0.4 | 0.85 ± 0.4 | 1.43 ± 0.7* | 1.66 ± 1.6*† |
| Percent WAT:body mass | 2.17 ± 1.0 | 2.95 ± 1.0 | 4.07 ± 1.6 | 4.18 ± 3.1 |
| Gastroc mass (mg) | 0.16 ± 0.03 | 0.17 ± 0.03 | 0.15 ± 0.02 | 0.15 ± 0.03 |
| Percent gastroc:body mass[a] | 0.56 ± 0.08 | 0.60 ± 0.08 | 0.40 ± 0.16 | 0.44 ± 0.08 |
| Soleus mass (mg) | 0.01 ± 0.0 | 0.01 ± 0.0 | 0.06 ± 0.2 | 0.01 ± 0.0 |
| Percent soleus: body mass | 0.04 ± 0.0 | 0.04 ± 0.0 | 0.14 ± 0.6 | 0.03 ± 0.0 |
| Uterus mass (mg) | 0.12 ± 0.0 | 0.10 ± 0.0 | 0.12 ± 0.05 | 0.12 ± 0.05 |
| Percent uterus mass:body mass | 0.5 ± 0.1 | 0.37 ± 0.1 | 0.38 ± 0.1 | 0.4 ± 0.2 |
| Frailty Index | | | 0.23 ± 0.07 | 0.22 ± 0.08 |

Data are mean ± SD. WAT, white adipose tissue. [a] p<0.05 main effect of age, * P <0.05 vs. young *APOE3*, † P<0.05 vs. young *APOE4*

**Supplemental Table 3.** Posterior Cerebral Artery characteristics

| **Variable** | **Young *APOE3*** | **Young *APOE4*** | **Old *APOE*3** | **Old *APOE4*** |
|---|---|---|---|---|
| Maximal Diameter (µm) | 141 ± 14 | 145 ± 11 | 132 ± 22 | 147 ± 18 |
| EC50, log M | | | | |
| ACh | -7.0 ± 0.4 | -7.1 ± 0.4 | -6.9 ± 0.4 | -7.1 ± 0.3 |
| Insulin | 1.6 ± 0.5 | 1.4 ±0.4 | 1.6 ± 0.5 | 1.6 ± 0.5 |
| ET1 | -7.2 ± 0.8 | -7.2 ± 0.7 | -7.5 ± 0.5 | -7.3 ± 0.7 |
| KCl | -7.6 ± 1.6 | -8.1 ± 0.5 | -8.3 ± 0.3 | -9.5 ± 0.3 |
| SNP | -7.1 ± 0.6 | -7.0 ± 0.5 | -7.5 ± 0.5 | -7.1 ± 0.6 |
| Preconstriction (%) | | | | |
| ACh | 37 ± 9 | 38 ± 13 | 35 ± 8 | 40 ± 10 |
| Insulin | 41 ± 10 | 38 ± 14 | 36 ± 11 | 40 ± 9 |
| SNP | 45 ± 14 | 50 ± 12 | 46 ± 14 | 49 ± 12 |

Data are mean ± SD. ACh, acetylcholine; ET1, endothelin-1; KCl, potassium chloride; SNP, sodium nitroprusside. No significant effects for 2x2 ANOVA analysis

**Supplemental Figure 1**

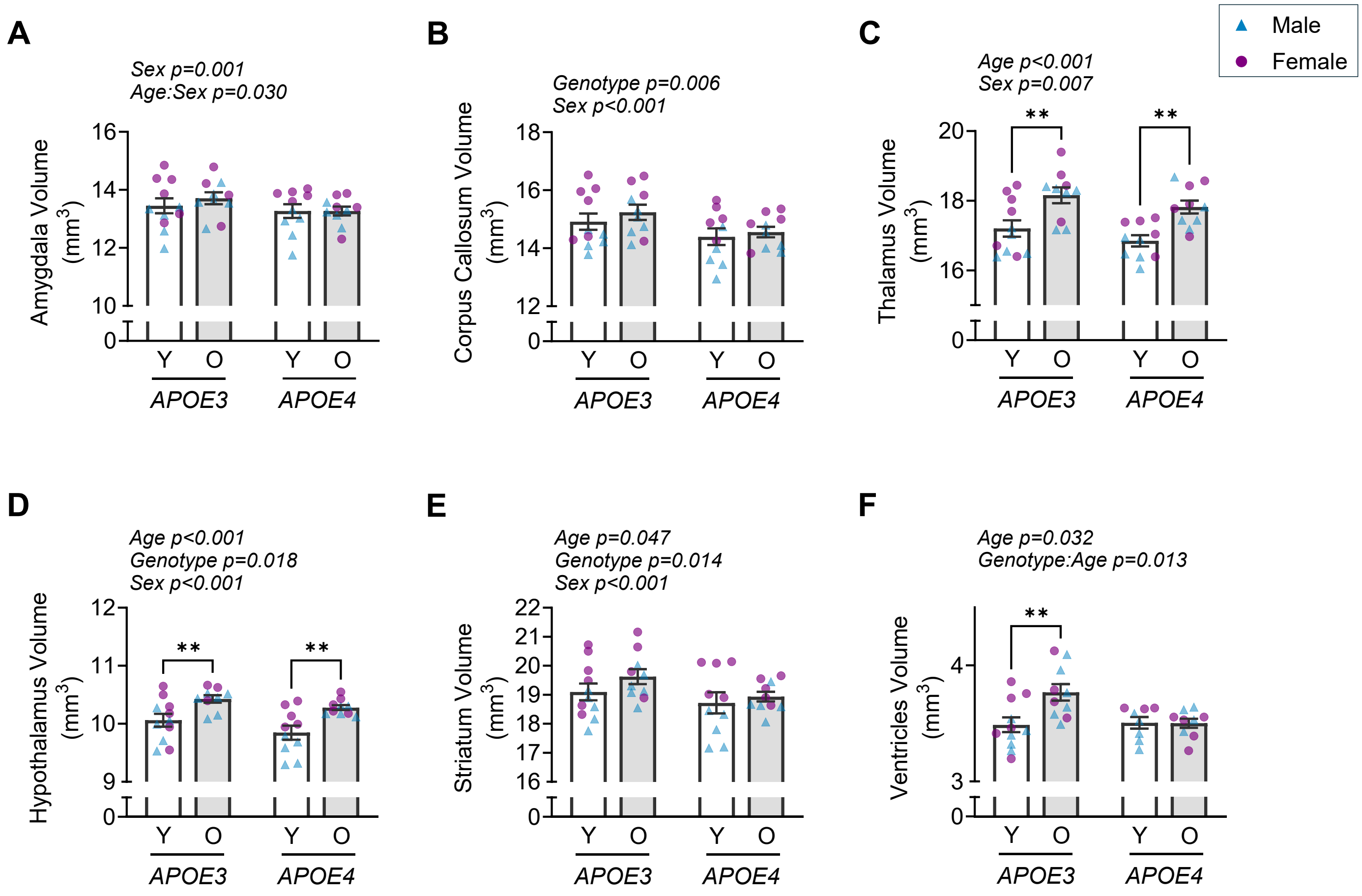


**Supplemental Figure 1. Effects of *APOE* genotype, age, and sex on regional brain volume.** From *in vivo* T2-weighted MRIs, volume of the (**A**) amygdala, (**B**) corpus callosum, (**C**) thalamus, (**D**) hypothalamus, (**E**) striatum, and (**F**) ventricles. Data analyzed by a three-way ANOVA with Tukey's multiple comparisons. Y, young (~6 months), O, old (~24 months). **p<0.01. n=4-6/group. Data are mean ± SEM.

**Supplemental Figure 2**

**Western Blot- Stain Free Blot**

Hippocampus- $ET_A$

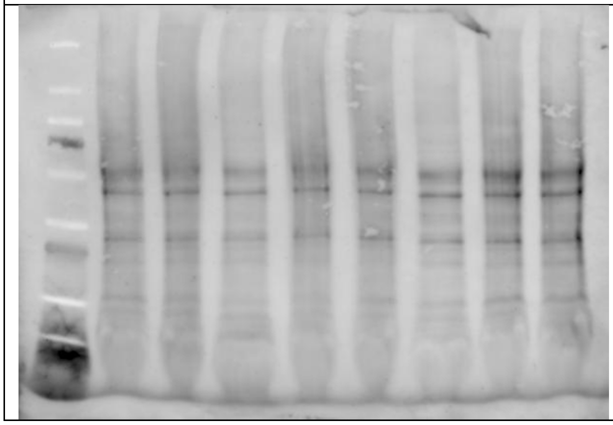

Hippocampus- $ET_B$, APOE

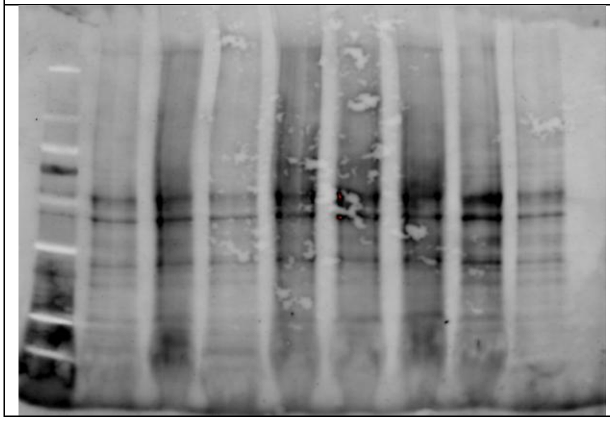

Aorta- $ET_1$

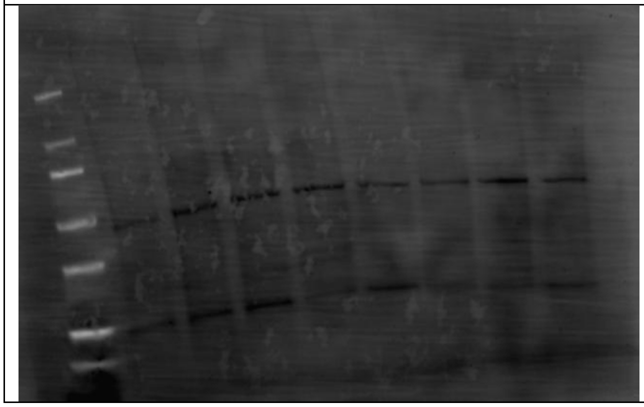
